# A minimal growth medium for *Pseudomonas azotoformans* associated with insect gut

**DOI:** 10.1101/2022.03.14.482786

**Authors:** Ismaela Cavallin, Bessem Chouaia

## Abstract

*Pseudomonas* spp. can be found associated with several insect hosts. Some of these strains play a crucial role in helping their hosts overcome biotic challenges such as plant defence effectors. Defined media and optimally minimal media are instrumental in studying the physiology and metabolic pathways of microorganisms. In the case of the insect-associated *Pseudomonas azotoformans* isolated from *Galeruca laticollis*’ gut, we tested six previously designed defined/minimal media. To determine the optimal medium, we compared the bacterium’s growth performances on the different defined/minimal media to its growth performance on a rich medium using the Dynamic Time Warping Algorithm. The growth results at both 25°C and 30°C indicated the SMM as the optimal medium, with bacterial growth performances being the closest to a rich medium. Therefore, this medium would be highly suitable to study the catabolic pathways for the degradation of some xenobiotics in *P. azotoformans*.

## Introduction

Bacteria of the genus *Pseudomonas* can be found in many habitats, including host-associated ones [1]. Several *Pseudomonas* strains have been found in plants [2–4] and animals [5–7] with a variety of phenotypes ranging from pathogenicity [3,8] to mutualism [5,9,10]. During recent years, some *Pseudomonas* strains were discovered to play a crucial role in helping different insects overcome biotic challenges such as pesticides or plant defence effectors [5,9,11]. Although the bacterium’s role in these symbioses is established, the molecular and biochemical mechanisms by which the symbionts are able to protect their respective hosts remain unknown.

Understanding how *Pseudomonas* species interact with their environment, host-associated or not, is of interest from a fundamental and applied perspective. For example, understanding how these bacteria are able to degrade a wide range of xenobiotics (such as pesticides) could help develop more ecologically-friendly approaches to deal with contaminations due to these xenobiotics. Since such metabolic investigations would ideally be carried out outside of the insect host, the development of suitable bacterial growth media that support good growth and produce consistently high viable cell counts is crucial. Such media are instrumental in studying certain aspects of bacterial metabolism at the biochemical and molecular level (i.e. transcriptomics).

However, for physiological and metabolical studies of these strains, defined/minimal media (liquid and agar) that support good growth and gives consistently high viable cell counts are a staple to unveil the mechanisms of action of metabolical processes and clarify the effects of changes in medium composition on cell growth and product formation. Defined media are culture media whose chemical composition is well known. A defined medium with only the ingredients necessary for the growth of a microorganism is called a minimal medium. Nevertheless, the number of ingredients that compose a minimal medium varies enormously and depends mainly on which microorganism is cultivated. Chemically defined minimal media allow for a finer metabolic profiling of the microorganism [12,13].

Although several defined or minimal nutritional media have been used for the genus *Pseudomonas*, most of the media were developed for the most common species (e.g., *P. syringae, P. aeruginosa, P. fluorescens*;) [14–16] and the growth performance of other strains on these media have rarely been investigated. Therefore, there is a need to test the already available minimal defined media and possibly develop more suitable ones for other *Pseudomonas* species.

Here, we aimed to determine an ideal minimal defined medium to grow a strain of *P. azotoformans*, isolated from an insect gut, in the laboratory. To this end, we compared the growth performance of this strain on several media to a rich medium (i.e. Luria Bertani broth).

## Materials and methods

### Strain

The bacterial strain was isolated from the gut of the beetle *Galeruca laticollis* plated on LB agar media and incubated at 30°C. The strain was initially identified based on its 16S rRNA gene sequence, and the taxonomic assignment was later confirmed through a phylogenomic analysis (data subject of a separate publication). The strain was identified as *Pseudomonas azotoformans* strain GL93.

### Culture media and growth conditions

In addition to LB media, which was used as a growth reference, five minimal media (namely SMM, M9, MOPS, MMV, and MMP) were tested (see supplementary file S1 for details). During the first test, bacteria grown on MMP produced fluorescent siderophores that could potentially interfere with the optical density (OD) readings. In order to minimize the production of these siderophores, the medium was supplemented with calcium chloride (CaCl_2_; supplementary figure S1).

The bacterium was incubated overnight at 30°C in a fresh LB medium prior to the experiments. Log-phase cells were collected by centrifugation at 2000 g for 10 minutes and washed three times in a sterile NaCl 9‰ solution to remove any remnants of the previous growth medium. Finally, the cells were concentrated to the desired concentration (10^8^ cells/ml), and 1 ml was used for the inoculation of different growth media. To test if the bacterium could grow on the different media, 100 ml of each medium were inoculated with 1 ml of a NaCl 9‰ solution containing 10^8^ bacterial cells. Bacterial growth at 30°C was monitored for seven days, taking OD readings at 24h intervals using an ONDA V-10 plus spectrophotometer.

Three minimal media (SMM, M9 and MMP+CaCl_2_) as well as LB proved suitable to grow the *Pseudomonas* strain (i.e. OD_medium_ > OD_LB_*0.2 after seven days) and were used in a second experiment. A 24-well plate was prepared with 200 µl of a given medium per well, one medium per row (see supplementary figure S2 for an illustration of the experimental design). Five out of the six wells for each medium were inoculated with 10 µl of the bacterial inoculate prepared as outlined above. The last well was used as blank. Bacterial growth was monitored for six days on a BioTek Synergy H1 multi-plate reader with OD readings at 600 nm every 15 minutes, preceded by 5 seconds of orbital shaking to homogenize the cultures. The experiment was carried out twice, once at 30°C and a second time at 25 °C.

### Data analyses

the R (https://cran.r-project.org) package growthcurver (https://github.com/sprouffske/growthcurver [17]) was used to infer the different terms of each logistic curve equation 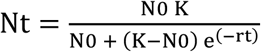, where Nt is the number of cells (or the absorbance reading) at time t, N0 is the initial cell count (or absorbance reading), K is the carrying capacity, and r is the growth rate. The Dynamic Time Warping (DTW; [18]) Algorithm from the package SimilarityMeasures (https://cran.r-project.org/web/packages/SimilarityMeasures/) was then used to determine the similarity between the different growth curves. First, we compared the growth curves between the replicates of each growth medium to exclude outliers. To this end, the values generated by pairwise comparisons of all growth curves were aggregated in a similarity distance matrix and visualized after hierarchical clustering. Secondly, we compared the averaged growth curves (i.e., curve resulting from the averaged values of the single curves) of the different media using the same approach used for the intra-medium comparisons.

### Metabolic pathway assessment

The metabolic capacities of P. azotoformans were checked using the four complete genomes available on the NCBI database (accession numbers: LT629702, CP041236, CP014546 and CP019856). In particular, the protein sequences of each genome were downloaded and submitted to GhostKOALA (ver 2.2, [19]) for metabolic pathway reconstruction. The pathways related to the central metabolism (e.g., glycolysis, pentose phosphate, TCA cycle, purine and pyrimidine synthesis, amino acid synthesis) were checked visually for completeness.

## Results and discussion

### Growth performance of *P. azotoformans* GL93 on different media

*P. azotoformans* was able to grow on all tested media but with very different growth performances (Table 1, Supplementary figure S3). As such, the bacterium reached the stationary phase on LB medium in less than 24h (mean = 18h), but it took longer to reach this phase on other media. It took ca 24h for the bacteria to reach the stationary phase on MMV, ca 36h on M9 and more than 48h on the other media. MOPS was the medium with the longest lag phase (mean = 36h). Similarly, the shortest generation time was observed on LB (ca. 1h) and the longest on both MMP and SMM (ca. 20h). However, *P. azotoformans* reached higher bacterial densities in MMP (OD_600_ = 1.18 ± 0.04) and SMM (OD_600_ = 1.3 ± 0.01) than in all other media, including LB (OD_600_ ≈ 1 ± 0.01). In the other media, the maximum value of OD_600_ remained inferior to one (namely M9: 0.3 ± 0.01, MMP + CaCl_2_: 0.2 ± 0.004, MMW: 0.12 ± 0.005, and MOPS: 0.03 ± 0.004). It is noteworthy that the addition of CaCl_2_ to the MMP medium reduced the production of siderophores (see Supplementary figure S1), but also had a notable effect on both the optical density (which decreased from an OD_600_ of 1.18 ± 0.04 to 0.2 ± 0.004) and on the generation time, lowering it from 20h to 12h. A possible explanation could be that, since the bacterium no longer produced siderophores, more resources could be invested into growth, resulting in a shortening of the time needed for cellular division. Considering the low bacterial density achieved on MMW and MOPS compared to LB (OD_600_ of 0.12 and 0.03, respectively), these two media were not included in subsequent experiments. The low growth performance on these two media could be due to a chemical being present at a highly limiting concentration or a growth-inhibiting factor for our strain.

**Table 1:** Optical density (OD) readings at 600 nm for each medium.

| Medium\time (h) | 0 | 24 | 48 | 72 | 144 | 168 |
| --- | --- | --- | --- | --- | --- | --- |
| LB | $0.03 \pm 0.01$ | $1.19 \pm 0.04$ | $0.98 \pm 0.04$ | $0.91 \pm 0.02$ | $0.97 \pm 0.04$ | $0.94 \pm 0.02$ |
| SMM | $0.02 \pm 0.02$ | $0.37 \pm 0.01$ | $0.53 \pm 0.02$ | $0.66 \pm 0.02$ | $1.16 \pm 0.01$ | $1.3 \pm 0.01$ |
| M9 | $0.02 \pm 0.02$ | $0.07 \pm 0.01$ | $0.42 \pm 0.02$ | $0.39 \pm 0.03$ | $0.24 \pm 0.01$ | $0.3 \pm 0.01$ |
| MOPS | 0 | $0.02 \pm 0$ | $0.03 \pm 0.01$ | $0.04 \pm 0.01$ | $0.03 \pm 0$ | $0.03 \pm 0$ |
| MMW | $0 \pm 0$ | $0.16 \pm 0$ | $0.18 \pm 0$ | $0.15 \pm 0$ | $0.1 \pm 0$ | $0.12 \pm 0$ |
| MMP | 0 | $0.32 \pm 0.01$ | $0.49 \pm 0.02$ | $0.65 \pm 0.02$ | $1.08 \pm 0.03$ | $1.18 \pm 0.04$ |
| MMP + $CaCl_2$ | $0 \pm 0$ | $0.12 \pm 0$ | $0.18 \pm 0$ | $0.2 \pm 0.01$ | $0.21 \pm 0.01$ | $0.2 \pm 0$ |

The reported values are the average of three readings and the standard error

### Identification of the best minimal medium for *P. azotoformans*

The growth of *P. azotoformans* on the selected media (SMM, M9, and MMP + CaCl2, hereafter MMP) was monitored every 15 minutes at two constant temperatures (i.e., 25°C and 30°C). The data obtained from 5 replicate cultures for each medium at both temperatures was used to infer the theoretical growth curve for each culture (Figs 1, 3). Comparing the growth curves of the five replicate cultures for each medium at both temperatures (Fig. 2, 4) revealed a certain degree of variability between replicates for some conditions. In particular, there was some variability in the exponential growth phase for M9 at both temperatures and for MMP at 30°C (Fig. 2, 4).

**Figure 1:**
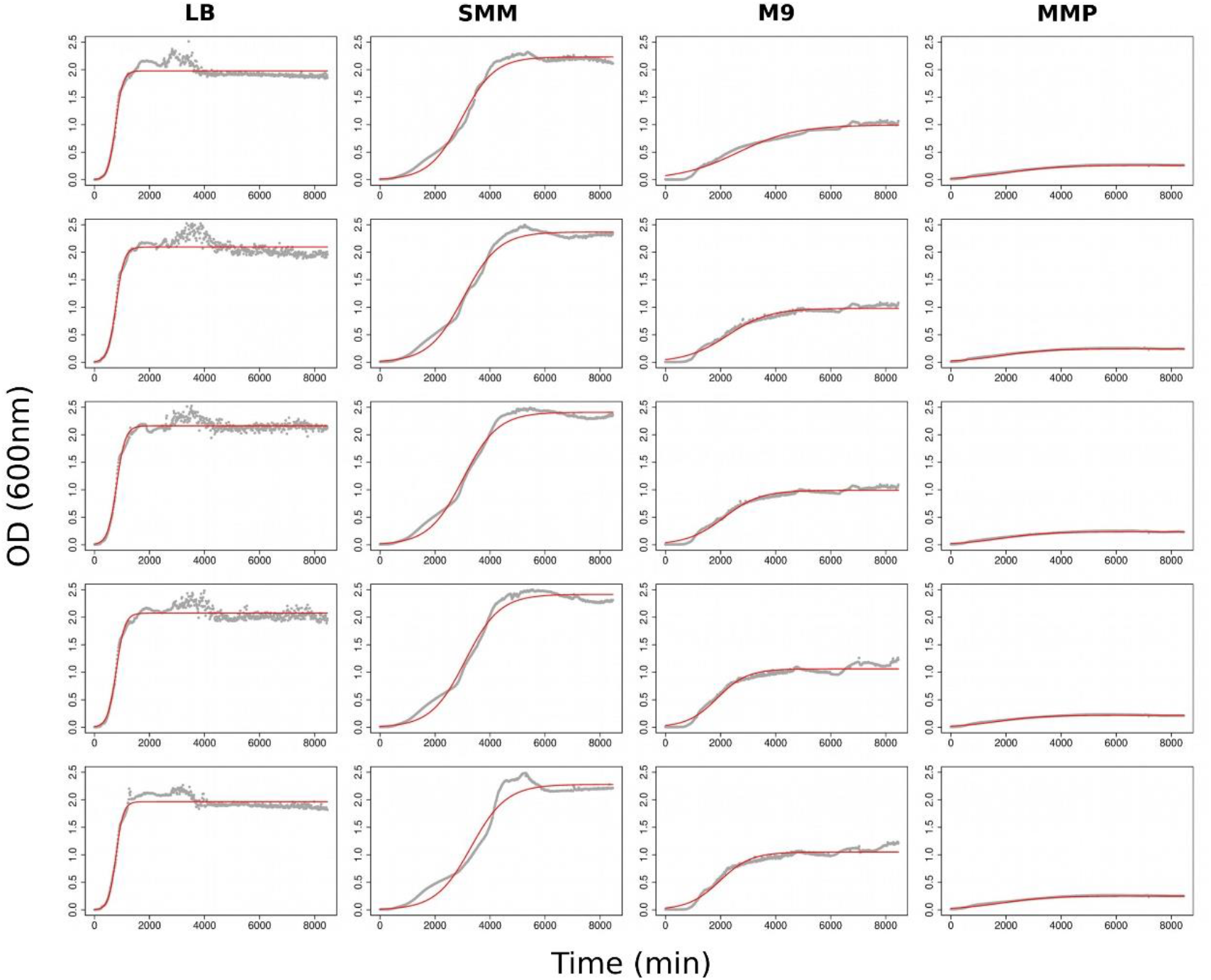
Growth curves obtained from the absorbance readings of the different media incubated at 25°C. The experimental curves (grey dots) correspond to the absorbance values recorded every 15 minutes for seven days. Each column corresponds to a medium, and each graph shows the growth performance in a single well (N=5 per medium). The red line represents the theoretical curve fitted over the experimental points.

**Figure 2:**
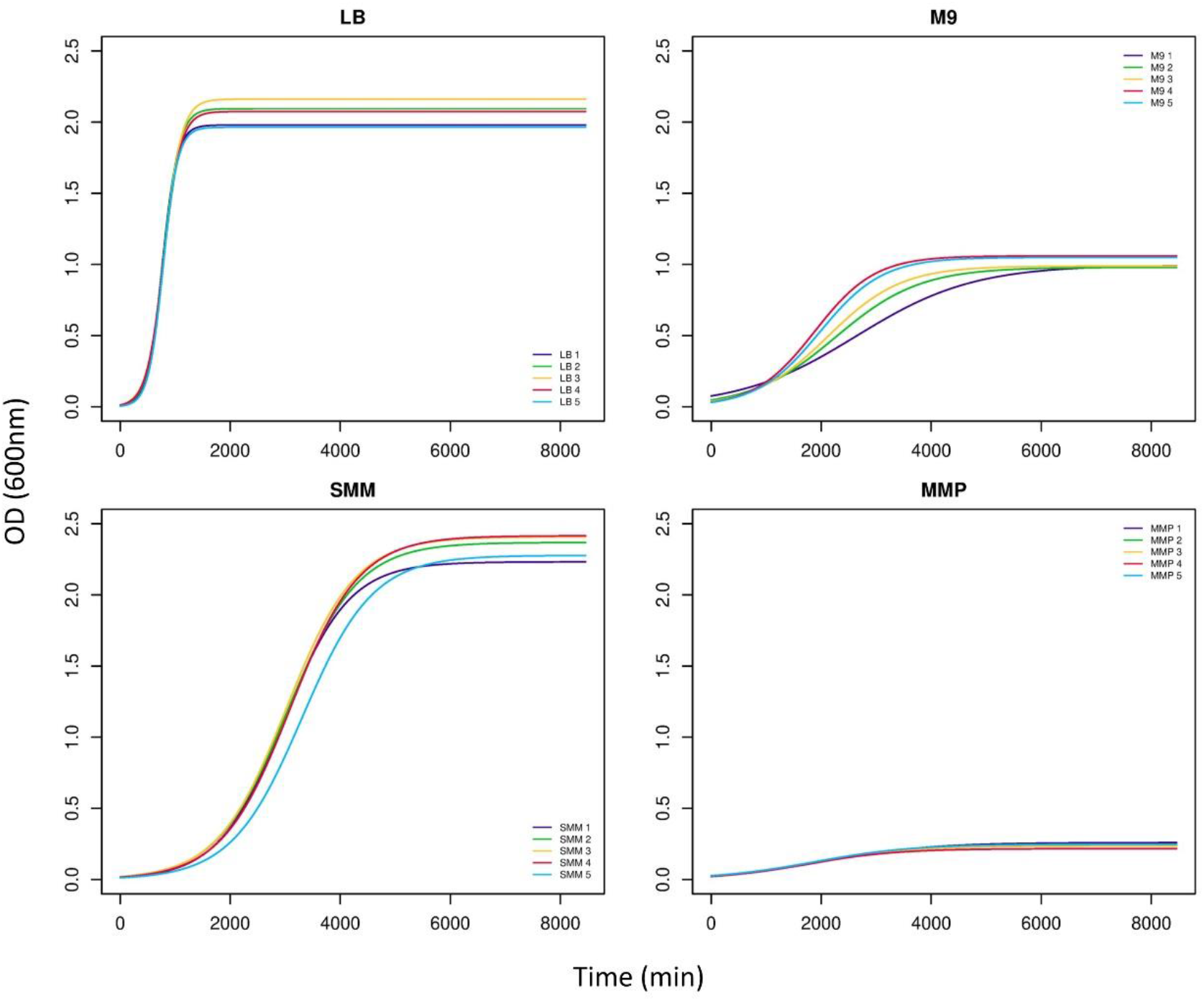
The theoretical growth curves of the 5 replicate cultures per medium at 25°C.

**Figure 3:**
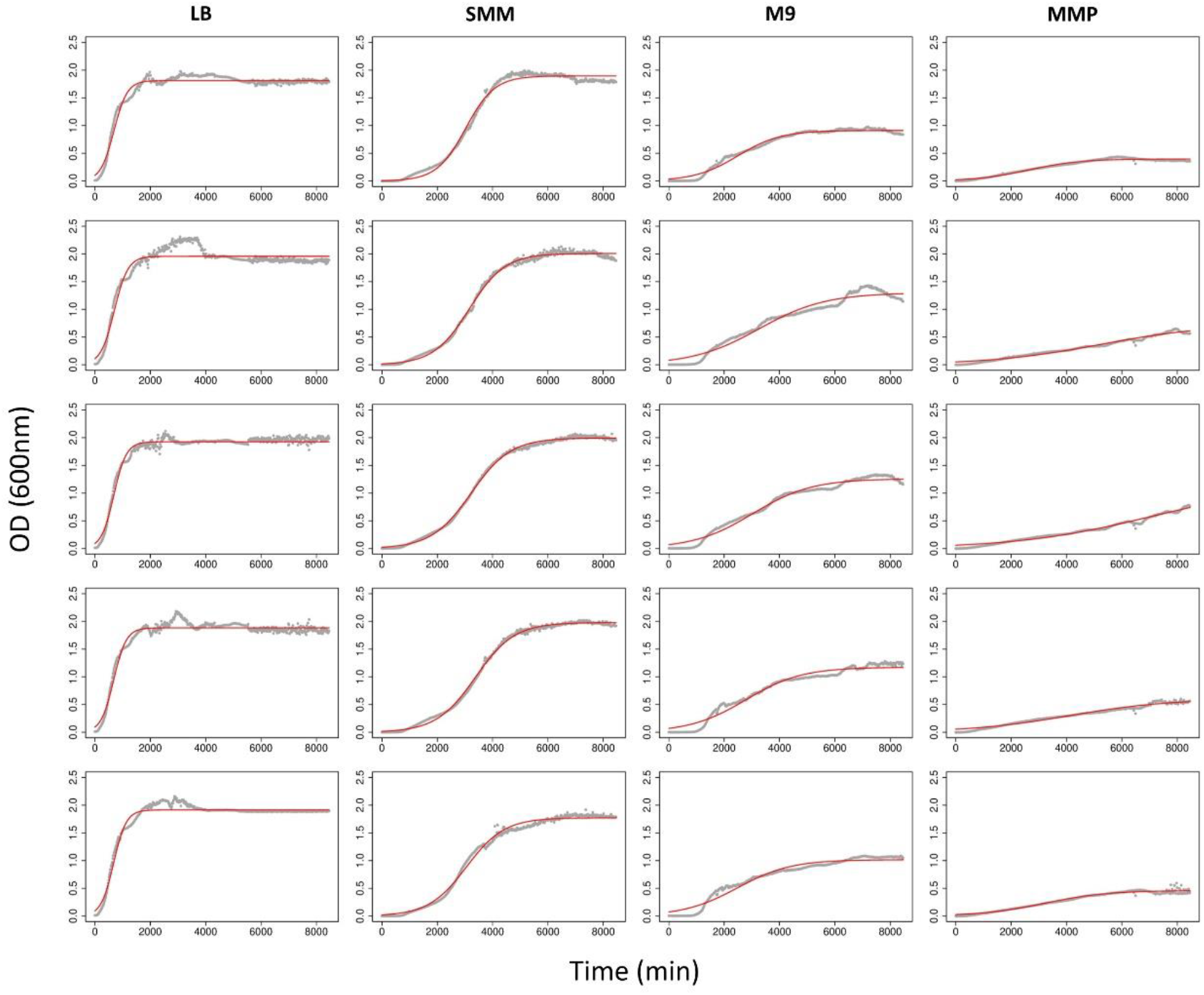
Growth curves obtained from the absorbance readings of the different media incubated at 30°C. The experimental curves (grey dots) correspond to the absorbance values recorded every 15 minutes for seven days. Each column corresponds to a medium, and each graph describes the growth performance in a single well (N=5 per medium). The red line represents the theoretical curve fitted over the experimental points.

**Figure 4:**
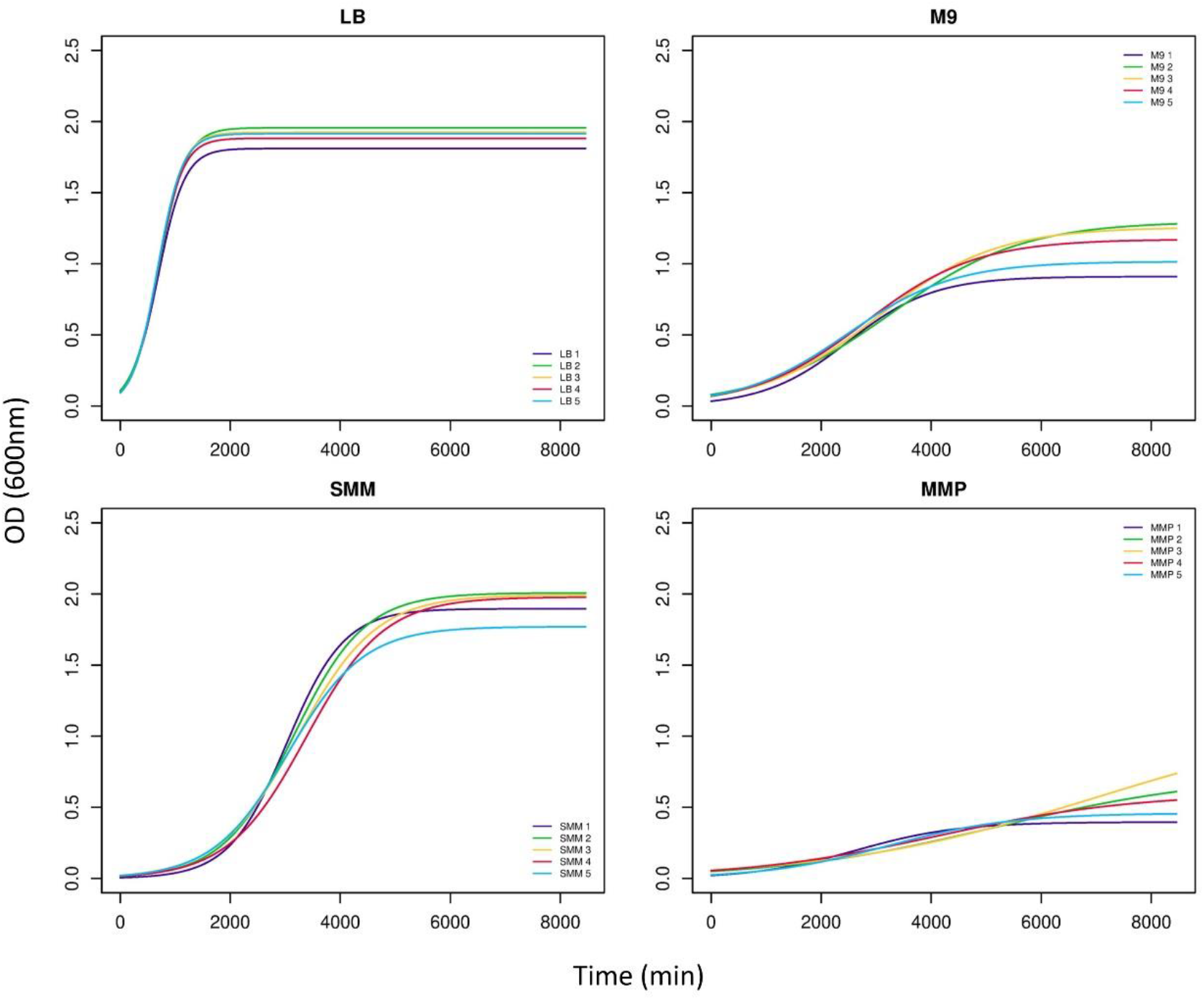
The theoretical growth curves of the 5 replicate cultures per medium at 30°C.

In order to determine which replicates represented significant deviations from the general trend of each medium, we applied the DTW algorithm on a pairwise distance matrix between the replicates. Using this approach, several replicates were indeed determined to be outliers: 1 LB replicate, 1 SMM replicate and 2 M9 replicates at 25°C (Supplementary Figure S4), as well as 1 SMM and 1 M9 replicate at 30°C (Supplementary Figure S6). No outliers were detected among the different replicates of MMP at both temperatures. The replicates determined to be outliers were removed and the remaining replicates were used to calculate the average theoretical growth curve for each medium at both temperatures (Supplementary Figures S5, S7).

Comparing the average growth curves of the three minimal media to that of the rich LB medium revealed that the SMM medium performed best and even surpassed the LB medium in terms of bacterial density (i.e. higher OD_600_) in the stationary phase (Fig. 5A, C). In contrast, the other two minimal media M9 and MMP could not sustain the same growth and vitality performance (Fig. 5). MMP was the most different from bacterial growth on the LB medium. This was also evident from the inferred generation time at each medium based on the average absorbance values: generation time at 25°C was 99 min, 432 min, 560 min and 609 min on LB, SMM, M9 and MMP, respectively (Supplementary figure S5; tables 2, 3).

**Figure 5:**
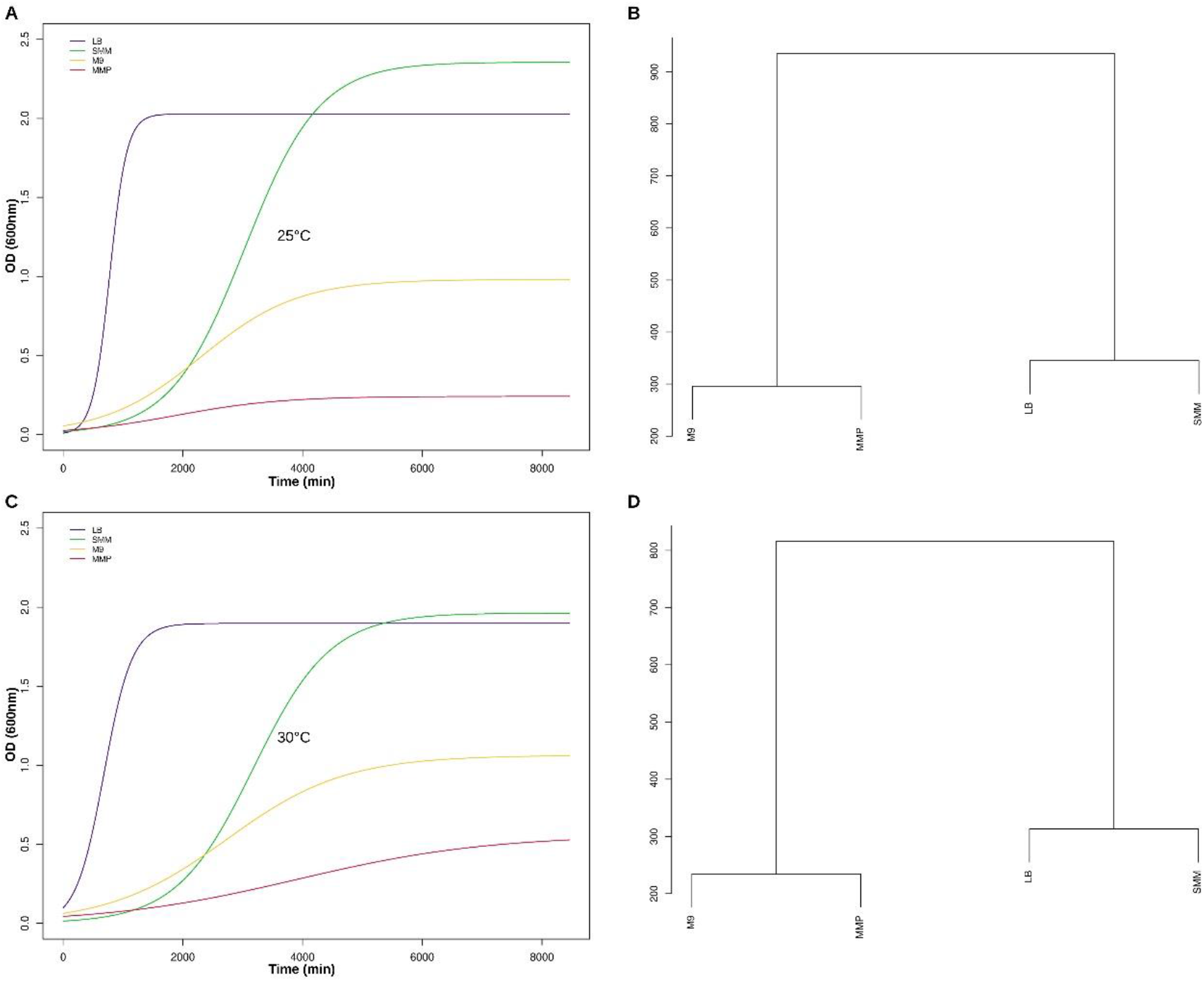
Comparison of the growth performance of *P. azotoformans* on the different media at 25°C (A, B) and 30°C (C, D). A and C: Averaged theoretical growth curves on the different media. B and D: Dendrograms of the curves’ similarity using the DTW metric.

**Table 2:** Logistic curve parameters estimated for each sample both at 30°C and 25°C.

| Sample | 30°C |  |  |  | 25°C |  |  |  |
| --- | --- | --- | --- | --- | --- | --- | --- | --- |
|  | K | N0 | r | Generation time | K | N0 | r | Generation time |
| LB1 | 1.811 | 0.101 | 0.004 | 168 | 1.979 | 0.007 | 0.007 | 93 |
| LB2 | 1.956 | 0.109 | 0.004 | 171 | 2.094 | 0.011 | 0.007 | 103 |
| LB3 | 1.924 | 0.091 | 0.004 | 157 | 2.161 | 0.014 | 0.006 | 110 |
| LB4 | 1.882 | 0.094 | 0.004 | 159 | 2.075 | 0.013 | 0.006 | 107 |
| LB5 | 1.914 | 0.094 | 0.004 | 158 | 1.964 | 0.006 | 0.007 | 93 |
| SMM1 | 1.896 | 0.006 | 0.002 | 363 | 2.232 | 0.015 | 0.002 | 412 |
| SMM2 | 2.008 | 0.015 | 0.001 | 448 | 2.368 | 0.02 | 0.002 | 444 |
| SMM3 | 1.994 | 0.022 | 0.001 | 497 | 2.408 | 0.02 | 0.002 | 436 |
| SMM4 | 1.978 | 0.017 | 0.001 | 490 | 2.415 | 0.017 | 0.002 | 433 |
| SMM5 | 1.770 | 0.020 | 0.001 | 474 | 2.277 | 0.013 | 0.002 | 445 |
| M91 | 0.911 | 0.035 | 0.001 | 536 | 0.996 | 0.076 | 0.001 | 736 |
| M92 | 1.298 | 0.080 | 0.001 | 831 | 0.98 | 0.048 | 0.001 | 530 |
| M93 | 1.257 | 0.069 | 0.001 | 739 | 0.989 | 0.039 | 0.002 | 462 |
| M94 | 1.172 | 0.070 | 0.001 | 700 | 1.061 | 0.033 | 0.002 | 379 |
| M95 | 1.016 | 0.072 | 0.001 | 670 | 1.05 | 0.032 | 0.002 | 395 |
| MMP1* | 0.397 | 0.022 | 0.001 | 643 | 0.261 | 0.027 | 0.001 | 663 |
| MMP2* | 0.733 | 0.050 | 0.001 | 1391 | 0.243 | 0.026 | 0.001 | 599 |
| MMP3* | 1.144 | 0.059 | 0.001 | 1670 | 0.236 | 0.024 | 0.001 | 592 |
| MMP4* | 0.601 | 0.055 | 0.001 | 1247 | 0.218 | 0.023 | 0.001 | 561 |
| MMP5* | 0.459 | 0.025 | 0.001 | 784 | 0.25 | 0.028 | 0.001 | 626 |

\* The medium contains CaCl<sub>2</sub> at the concentration reported in the supplementary file

**Table 3:** Logistic curve parameters estimated for the averaged curve for each medium both at 30°C and 25°C.

| sample | 30°C |  |  |  | 25°C |  |  |  |
| --- | --- | --- | --- | --- | --- | --- | --- | --- |
|  | K | N0 | r | Generation time | K | N0 | r | Generation time |
| LB | 1.897 | 0.098 | 0.004 | 163 | 2.028 | 0.009 | 0.007 | 99 |
| SMM | 1.963 | 0.014 | 0.002 | 446 | 2.355 | 0.018 | 0.002 | 432 |
| M9 | 1.063 | 0.062 | 0.001 | 684 | 0.981 | 0.054 | 0.001 | 560 |
| MMP* | 0.557 | 0.043 | 0.001 | 1100 | 0.241 | 0.025 | 0.001 | 610 |

\* The medium contains CaCl<sub>2</sub> at the concentration reported in the supplementary file

When comparing the growth performances on the different media at 25° and 30°C, it emerged that *Pseudomonas* generally grew more slowly at 30°C than at 25°C. Hence, generation time at 30°C was 162 min, 446 min, 684 min and 1100 min on LB, SMM, M9 and MMP, respectively (Supplementary figure S7; tables 2, 3). SMM represents the exception to this trend, since bacteria growing on SMM had similar growth rates at both temperatures (average generation time at 25°C: 432 min *vs* average generation time at 30°C: 446). Despite growing at a similar rate, bacterial density remained lower at 30°C than at 25°C (Fig. 5 A.C).

Taken together, *P. azotoformans* grown on SMM performed best compared to the other minimal media (shorter generation time and higher bacterial density). However, it was not comparable to growth on a rich medium.

Based on the metabolism of *P. azotoformans* inferred from the available genomes on the NCBI database (Table 4), we can hypothesize that the higher growth performance on the SMM medium could be due to minimal composition. SMM contains the necessary glucose as a carbon source, glutamine as a nitrogen source, K_2_HPO_4_ as a phosphate source and MgSO_4_ as a sulfur source. These four ingredients are sufficient in *P. azotoformans* for the biosynthesis of all the needed building blocks for metabolism and growth. The addition of other nutrients, such as for the other media tested, will probably trigger other, possibly secondary, metabolic pathways. These active metabolic pathways could divert resources (e.g., ATP, metabolites) from the central metabolism, thus slowing the growth and multiplication of the bacterial cells. This may result in slower performances when *P. azotoformans* is incubated in more complex media.

**Table 4:**
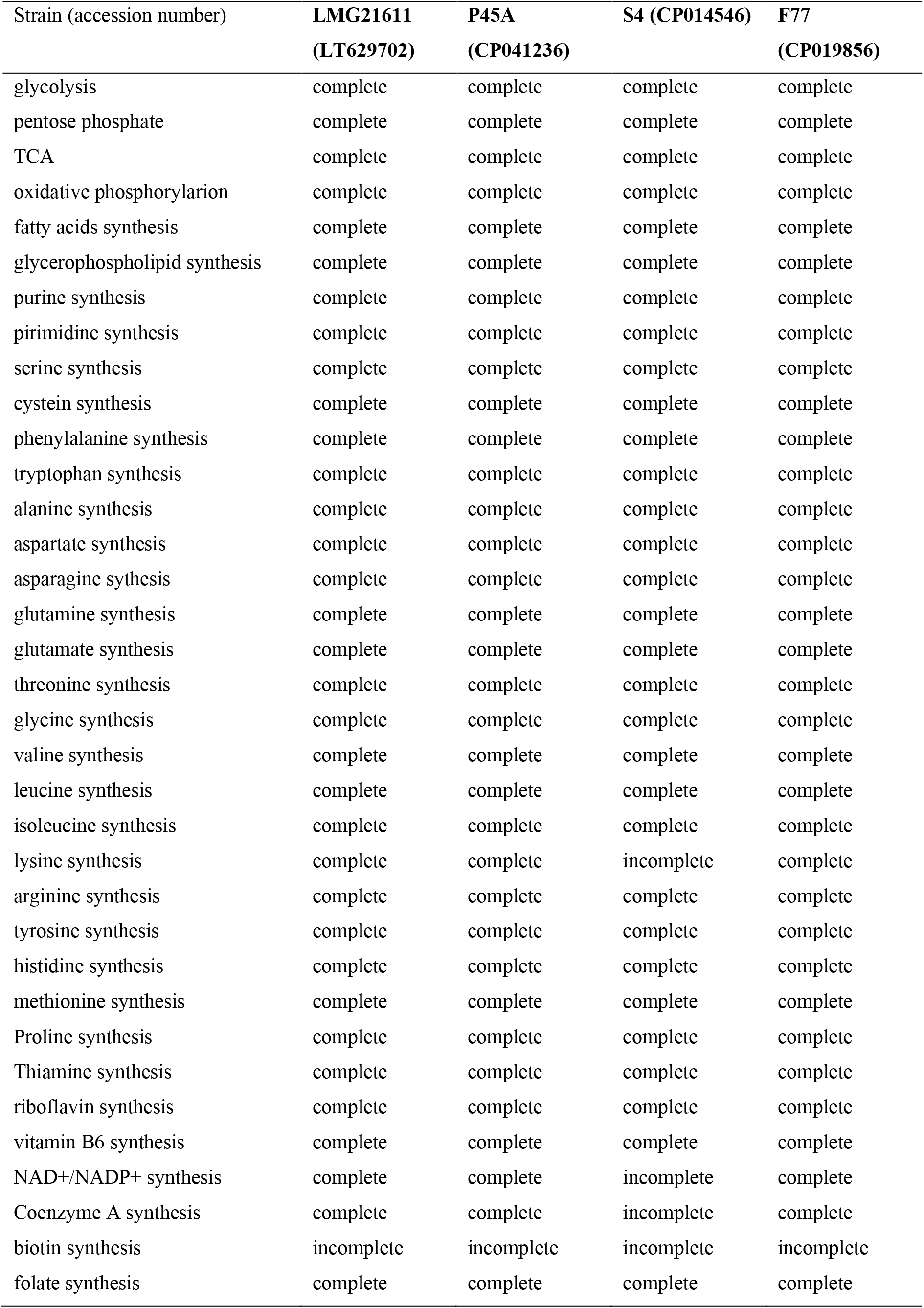
Level of completeness of the different metabolic pathways that are part of the central metabolism in four P. azotoformans strains.

| Strain (accession number) | LMG21611<br>(LT629702) | P45A<br>(CP041236) | S4 (CP014546) | F77<br>(CP019856) |
| --- | --- | --- | --- | --- |
| glycolysis | complete | complete | complete | complete |
| pentose phosphate | complete | complete | complete | complete |
| TCA | complete | complete | complete | complete |
| oxidative phosphorylation | complete | complete | complete | complete |
| fatty acids synthesis | complete | complete | complete | complete |
| glycerophospholipid synthesis | complete | complete | complete | complete |
| purine synthesis | complete | complete | complete | complete |
| pyrimidine synthesis | complete | complete | complete | complete |
| serine synthesis | complete | complete | complete | complete |
| cysteine synthesis | complete | complete | complete | complete |
| phenylalanine synthesis | complete | complete | complete | complete |
| tryptophan synthesis | complete | complete | complete | complete |
| alanine synthesis | complete | complete | complete | complete |
| aspartate synthesis | complete | complete | complete | complete |
| asparagine synthesis | complete | complete | complete | complete |
| glutamine synthesis | complete | complete | complete | complete |
| glutamate synthesis | complete | complete | complete | complete |
| threonine synthesis | complete | complete | complete | complete |
| glycine synthesis | complete | complete | complete | complete |
| valine synthesis | complete | complete | complete | complete |
| leucine synthesis | complete | complete | complete | complete |
| isoleucine synthesis | complete | complete | complete | complete |
| lysine synthesis | complete | complete | incomplete | complete |
| arginine synthesis | complete | complete | complete | complete |
| tyrosine synthesis | complete | complete | complete | complete |
| histidine synthesis | complete | complete | complete | complete |
| methionine synthesis | complete | complete | complete | complete |
| Proline synthesis | complete | complete | complete | complete |
| Thiamine synthesis | complete | complete | complete | complete |
| riboflavin synthesis | complete | complete | complete | complete |
| vitamin B6 synthesis | complete | complete | complete | complete |
| NAD<sup>+</sup>/NADP<sup>+</sup> synthesis | complete | complete | incomplete | complete |
| Coenzyme A synthesis | complete | complete | incomplete | complete |
| biotin synthesis | incomplete | incomplete | incomplete | incomplete |
| folate synthesis | complete | complete | complete | complete |

## Conclusion

An optimal medium allows for good microbial growth, which can be summarised as short generation time and high cell concentration in solution. In light of the results achieved here, the SMM medium is the most suitable medium for studying the metabolism of *P. azotoformans*. SMM outperformed the other two media (i.e., M9 and MMP), and the bacterial growth performances were consistently closer to those on LB. Furthermore, while the optimal growth temperature is 25°C, the SMM medium was the least susceptible to performance variations at a suboptimal growth temperature.

The better growth performance on the SMM medium could be partly explained by the fact that the medium contains the necessary nutrients sufficient for the biosynthesis of all the needed building blocks for metabolism and growth. In addition, *P. azotoformans* is a phylogenetic closer to P. fluorescens [20], which SMM was originally designed, than other *Pseudomonas* species. In contrast, most of the other minimal media tested here had been designed for *P. aeruginosa*, except for the more generic MMP. The phylogenetic proximity with *P. fluorescens* may suggest similar metabolic needs between the two species. Thus, a medium developed for one of the two species could offer better growth performances for the second species than media developed for more distant relatives.

## Acknowledgements

BC would like to thank Jessica Dittmer for the critical reading of the paper.

## Supplementary materials

**Supplementary file S1**: Composition of the different defined media used in this study.

**Supplementary figure S1**: The effect of adding CaCl_2_ to the MMP medium on the production of siderophores. A: *Pseudomonas* grown in MMP. The yellow colour indicates the production of siderophores. B: *Pseudomonas* grown in MMP + CaCl2. The transparent medium indicates that no siderophores were produced.

**Supplementary figure S2**: The experimental setup with the media disposition on the plate for the second part of the experiment (i.e., continuous long time monitoring using a plate reader).

**Supplementary figure S3**: Comparison of the growth performance of *P. azotoformans* on the six different media at 30°C. The curves represent the averaged theoretical growth curves for each medium.

**Supplementary figure S4**: Dendrograms showing the similarity between the theoretical growth curves of the five replicates for each medium at 25°C. The dotted red line delineates outliers based on the DTW metric (75% similarity threshold).

**Supplementary figure S5**: Averaged growth curve for each medium at 25°C after excluding outliers. The experimental curves (grey dots) correspond to the absorbance values recorded every 15 minutes for seven days. The red line represents the theoretical curve fitted over the experimental points. The inserts show the generation time inferred from the averaged absorbance values.

**Supplementary figure S6**: Dendrograms showing the similarity between the theoretical growth curves of the five replicates for each medium at 30°C. The dotted red line delineates outliers based on the DTW metric (75% similarity threshold).

**Supplementary figure S7**: Averaged growth curve for each medium at 30°C after excluding outliers. The experimental curves (grey dots) correspond to the absorbance values recorded every 15 minutes for seven days. The red line represents the theoretical curve fitted over the experimental points. The inserts show the generation time inferred from the averaged absorbance values.

## Notes

### Competing Interest Statement

The authors have declared no competing interest.

## References

1. Peix A, Ramírez-Bahena MH, Velázquez E. Historical evolution and current status of the taxonomy of genus Pseudomonas. Infect Genet Evol. 2009;9: 1132–1147. doi:10.1016/j.meegid.2009.08.001

2. Anderson AJ, Kim YC. Insights into plant-beneficial traits of probiotic Pseudomonas chlororaphis isolates. J Med Microbiol. 2020;69: 361–371. doi:10.1099/jmm.0.001157

3. Xin XF, Kvitko B, He SY. Pseudomonas syringae: What it takes to be a pathogen. Nat Rev Microbiol. 2018;16: 316–328. doi:10.1038/nrmicro.2018.17

4. Flury P, Vesga P, Dominguez-Ferreras A, Tinguely C, Ullrich CI, Kleespies RG, et al. Persistence of root-colonising Pseudomonas protegens in herbivorous insects throughout different developmental stages and dispersal to new host plants. ISME J. 2019;13: 860–872. doi:10.1038/s41396-018-0317-4

5. Shukla SP, Beran F. Gut microbiota degrades toxic isothiocyanates in a flea beetle pest. Mol Ecol. 2020;29: 4692–4705. doi:10.1111/mec.15657

6. Zhukova M, Sapountzis P, Schiøtt M, Boomsma JJ. Diversity and Transmission of Gut Bacteria in Atta and Acromyrmex Leaf-Cutting Ants during Development. Frontiers in Microbiology. 2017. p. 1942. doi:10.3389/fmicb.2017.01942

7. Montagna M, Gómez-Zurita J, Giorgi A, Epis S, Lozzia GC, Bandi C. Metamicrobiomics in herbivore beetles of the genus Cryptocephalus (Chrysomelidae): toward the understanding of ecological determinants in insect symbiosis. Insect Sci. 2015;22: 340–352. doi:10.1111/1744-7917.12143

8. Dolan SK, Kohlstedt M, Trigg S, Ramirez PV, Kaminski CF, Wittmann C, et al. Contextual flexibility in Pseudomonas aeruginosa central carbon metabolism during growth in single carbon sources. MBio. 2019;11: e02684–19. doi:10.1128/mBio.02684-19

9. Ceja-Navarro JA, Vega FE, Karaoz U, Hao Z, Jenkins S, Lim HC, et al. Gut microbiota mediate caffeine detoxification in the primary insect pest of coffee. Nat Commun. 2015;6: 7618. doi:10.1038/ncomms8618

10. Pieterse CMJ, Zamioudis C, Berendsen RL, Weller DM, Van Wees SCM, Bakker PAHM. Induced systemic resistance by beneficial microbes. Annu Rev Phytopathol. 2014;52: 347–375. doi:10.1146/annurev-phyto-082712-102340

11. Wang GH, Berdy BM, Velasquez O, Jovanovic N, Alkhalifa S, Minbiole KPC, et al. Changes in Microbiome Confer Multigenerational Host Resistance after Sub-toxic Pesticide Exposure. Cell Host Microbe. 2020;27: 213-224.e7. doi:10.1016/j.chom.2020.01.009

12. Singleton C, Gilman J, Rollit J, Zhang K, Parker DA, Love J. A design of experiments approach for the rapid formulation of a chemically defined medium for metabolic profiling of industrially important microbes. PLoS One. 2019;14: 7–11. doi:10.1371/journal.pone.0218208

13. Roth JR. Genetic Techniques in Studies of Bacterial Metabolism. Methods Enzymol. 1970;17: 3–35. doi:10.1016/0076-6879(71)17165-0

14. LaBauve AE, Wargo MJ. Growth and laboratory maintenance of Pseudomonas aeruginosa. Curr Protoc Microbiol. 2012; 1–8. doi:10.1002/9780471729259.mc06e01s25

15. Mishek HP, Stock SA, Florick JDE, Blomberg WR, Franke JD. Development of a chemically-defined minimal medium for studies on growth and protein uptake of Gemmata obscuriglobus. J Microbiol Methods. 2018;145: 40–46. doi:10.1016/j.mimet.2017.12.010

16. Sörensen M, Khakimov B, Nurjadi D, Boutin S, Yi B, Dalpke AH, et al. Comparative evaluation of the effect of different growth media on in vitro sensitivity to azithromycin in multi-drug resistant Pseudomonas aeruginosa isolated from cystic fibrosis patients. Antimicrob Resist Infect Control. 2020;9: 1–7. doi:10.1186/s13756-020-00859-7

17. Sprouffske K, Wagner A. Growthcurver: An R package for obtaining interpretable metrics from microbial growth curves. BMC Bioinformatics. 2016;17: 17–20. doi:10.1186/s12859-016-1016-7

18. Berndt DJ, Clifford J. Using dynamic time warping to find patterns in time series. Work Knowl Knowl Discov Databases. 1994;398: 359–370. Available: http://www.aaai.org/Papers/Workshops/1994/WS-94-03/WS94-03-031.pdf

19. Kanehisa M, Sato Y, Morishima K. BlastKOALA and GhostKOALA: KEGG tools for functional characterization of genome and metagenome sequences. J. Mol. Biol. 2016;428: 726–731. doi:10.1016/j.jmb.2015.11.006

20. Anzai Y, Kim H, Park J, Wakabayashi H, Oyaizu H. Phylogenetic affiliation of the pseudomonads based on 16S rRNA sequence. Int J Syst Evol Microbiol. 2000;50: 1563– 1589. doi: 10.1099/00207713-50-4-1563

