## Supplementary material and methods for "A minimal growth medium for *Pseudomonas azotoformans* associated with insect gut"

**Table S1: composition of the minimal medium MMW (Whitchurch et al, 2002)**

| compound | amount (mM) |
|---|---|
| Ammonium sulfate | 1.51 |
| Boric acid | 8.086e-05 |
| Calcium chloride anhydrous | 0.01 |
| Calcium sulfate | 0.001469 |
| Cobaltous sulfate | 3.557e-05 |
| Cupric sulfate | 0.0001253 |
| Dibasic sodium phosphate | 3.37 |
| Ferrous sulfate | 0.0007194 |
| Iron(III) chloride | 0.001 |
| Magnesium chloride | 5.0 |
| Manganese sulfate | 0.0001325 |
| Potassium dihydrogen phosphate | 2.2 |
| Sodium chloride | 179.0 |
| Sodium citrate | 0.1 |
| Sodium molybdate | 4.856e-05 |
| Zinc sulfate | 6.955e-05 |

**Table S2: M9 minimal salts medium (Miller, 1972)**

| compound | amount for 100 mL |
|---|---|
| M9 salts (5X) * | 20 mL |
| Glucose (20%) | 2 mL |
| $MgSO_4$ (1M) | 200 µL |
| $CaCl_2$ (1M) | 10 µL |
| $H_2O$ | 78 mL |

* M9 salts composition:

| compound | amount (g/L) |
|---|---|
| $Na_2HPO_4•7H_2O$ | 64 |
| $KH_2PO_4$ | 15 |
| NaCl | 2.5 |
| $NH_4Cl$ | 5.0 |

**Table S3: MOPS (Neidhardt et *al*., 1974; LaBauve and Wargo, 2012)**

| Component | Stock | Volume for 500 mL | Final concentration |
|---|---|---|---|
| 10X MOPS stock * | See recipe * | 50 mL | See recipe * |
| Deionized water | N/A | 400 mL | N/A |
| Glucose | 1 M | 10 mL | 20 mM |
| CaCl2 | 53 mM | 300 µL | 32 µM |
| K2SO4 | 27.5 mM | 5 mL | 0.29 mM |
| K2HPO4 | 172.8 mM | 5 mL | 1.32 mM |

* MOPS stock components (10X)

| compound | Stock | Volume for 500 mL | Concentration at 1X |
|---|---|---|---|
| MOPS | 1 M (pH 7.5) | 200 mL | 40 nM |
| Tricine | 1 M (pH 7.5) | 20 mL | 4 mM |
| $FeSO_4$ | 18.4 mM | 5 mL | 0.01 mM |
| $NH_4Cl$ | 1.9 M | 25 mL | 9.52 mM |
| $CaCl_2$ | 53 mM | 50 µL | 0.5 µM |
| $MgCl_2$ (hexahydrate) | 512 mM | 5 mL | 0.52 mM |
| NaCl | 5 M | 50 mL | 50 mM |
| Micronutrients stock ** | 100X | 5 mL | See recipe ** |

**Micronutrient stock for MOPS

| compound | Mg/100 mL | Stock concentration (µM) |
|---|---|---|
| Ammonium molybdate tetrahydrate | 0.3 | 3 |
| Boric acid | 2.4 | 400 |
| Cobalt chloride | 0.7 | 30 |
| Cupric sulfate | 0.3 | 10 |
| Manganese sulfate | 1.6 | 80 |
| Zinc sulfate | 0.3 | 10 |

**Table S4: modified SMM (Cheng et *al*., 1995)**

| compound | amount (g/L) |
|---|---|
| Glucose | 20 |
| L-glutamine | 2 |
| $K_2HPO_4$ | 1 |
| $MgSO_4 \cdot 7H_2O$ | 0.5 |

**Table S4: modified MMP (Wick, 2010)**

| compound | amount for 1 L |
|---|---|
| $H_2O$ | 985 mL |
| Glycerol | 15 mL |
| L-Glutamine | 5 g |
| $K_2HPO_4$ | 1.5 g |
| $MgSO_4$ | 0.2 g |
| $CaCl_2$ (1M) * | 10 µL |

* the initial medium did not contain $CaCl_2$, it was added later to minimise the production of siderophores by *P. azotoformans*

### References


Cheng C-M, Doyle MP, Luchansky JB. 1995. Identification of *Pseudomonas fluorescens* Strains Isolated from Raw Pork and Chicken That Produce Siderophores Antagonistic towards Foodborne Pathogens. J Food Prot. 58:1340-1344. doi: 10.4315/0362-028X-58.12.1340.

LaBauve AE, Wargo MJ. 2012. Growth and Laboratory Maintenance of *Pseudomonas aeruginosa*. Current Protocols in Microbiology. 25:6E.1.1-6E.1.8. doi:10.1002/9780471729259.mc06e01s25

Miller JH. 1972. Experiments in molecular genetics, Cold Spring Harbor Laboratory. doi: 10.1101/pdb.rec12295.

Neidhardt FC, Bloch PL, Smith DF. 1974. Culture medium for Enterobacteria. J. Bacteriol. 119:736-747.

Wick R. the Culture Media for Plant Pathogenic Fungi and Bacteria, University of Massachusetts. https://wiki.bugwood.org/Minimal_medium_(Pseudomonas). Last accessed 20/12/2020.

Whitchurch CB, Tolker-Nielsen T, Ragas PC, Mattick JS. 2002. Extracellular DNA required for bacterial biofilm formation. Science, 295:1487. doi: 10.1126/science.295.5559.1487.
