## Supplementary figures and images for "A minimal growth medium for *Pseudomonas azotoformans* associated with insect gut"

### Supplementary figure S1

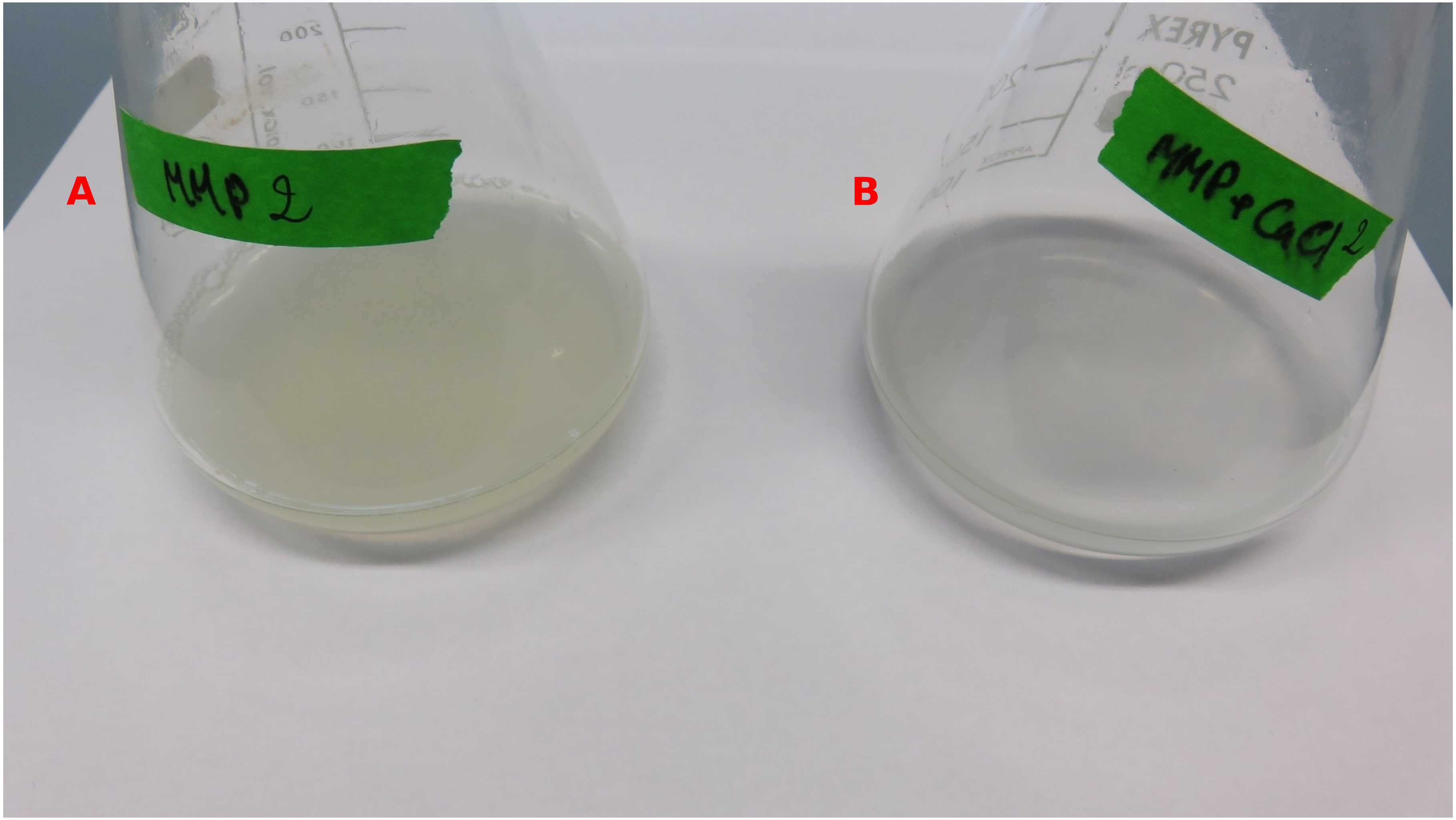

### Supplementary figure S3

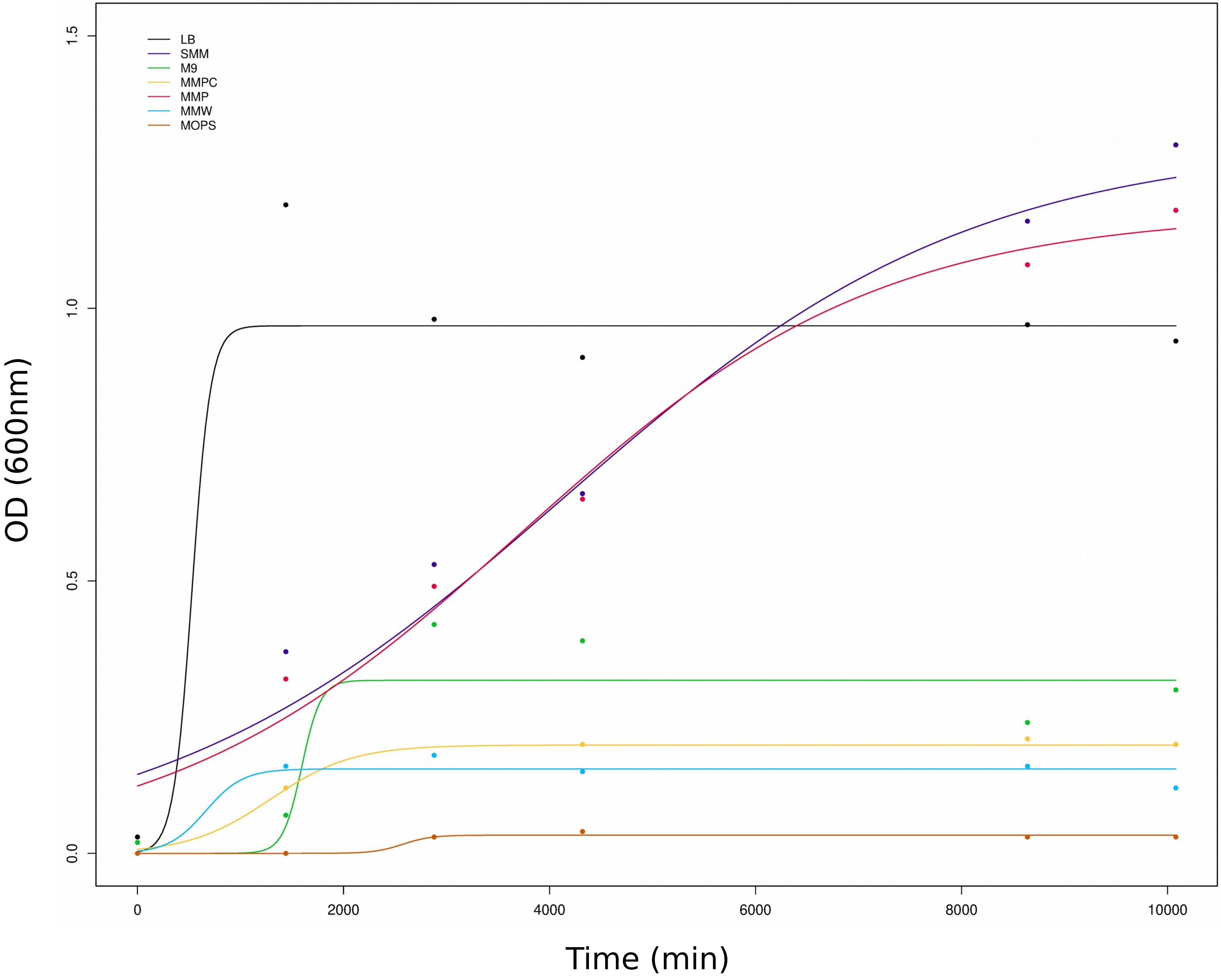

### Supplementary figure S4

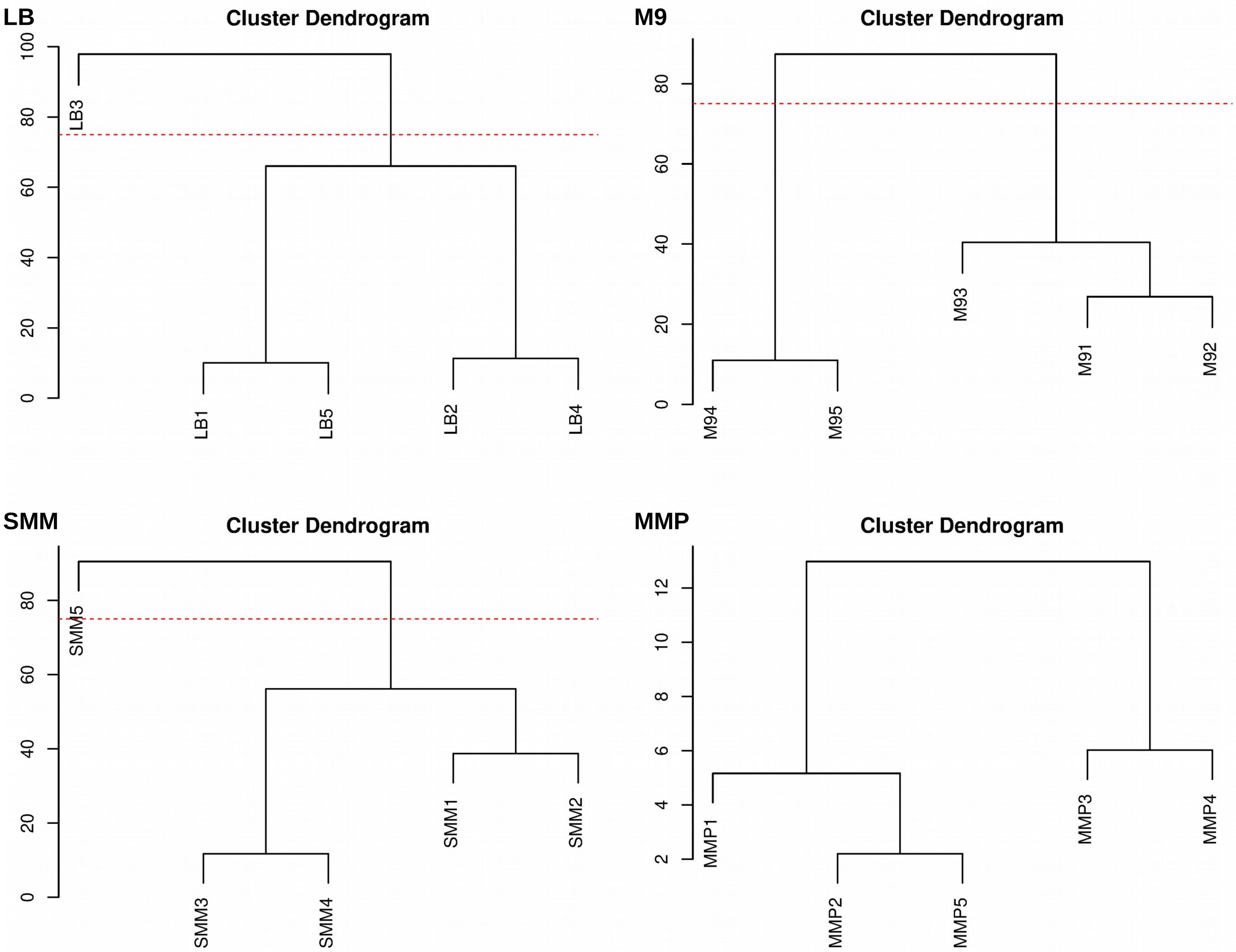

### Supplementary figure S5

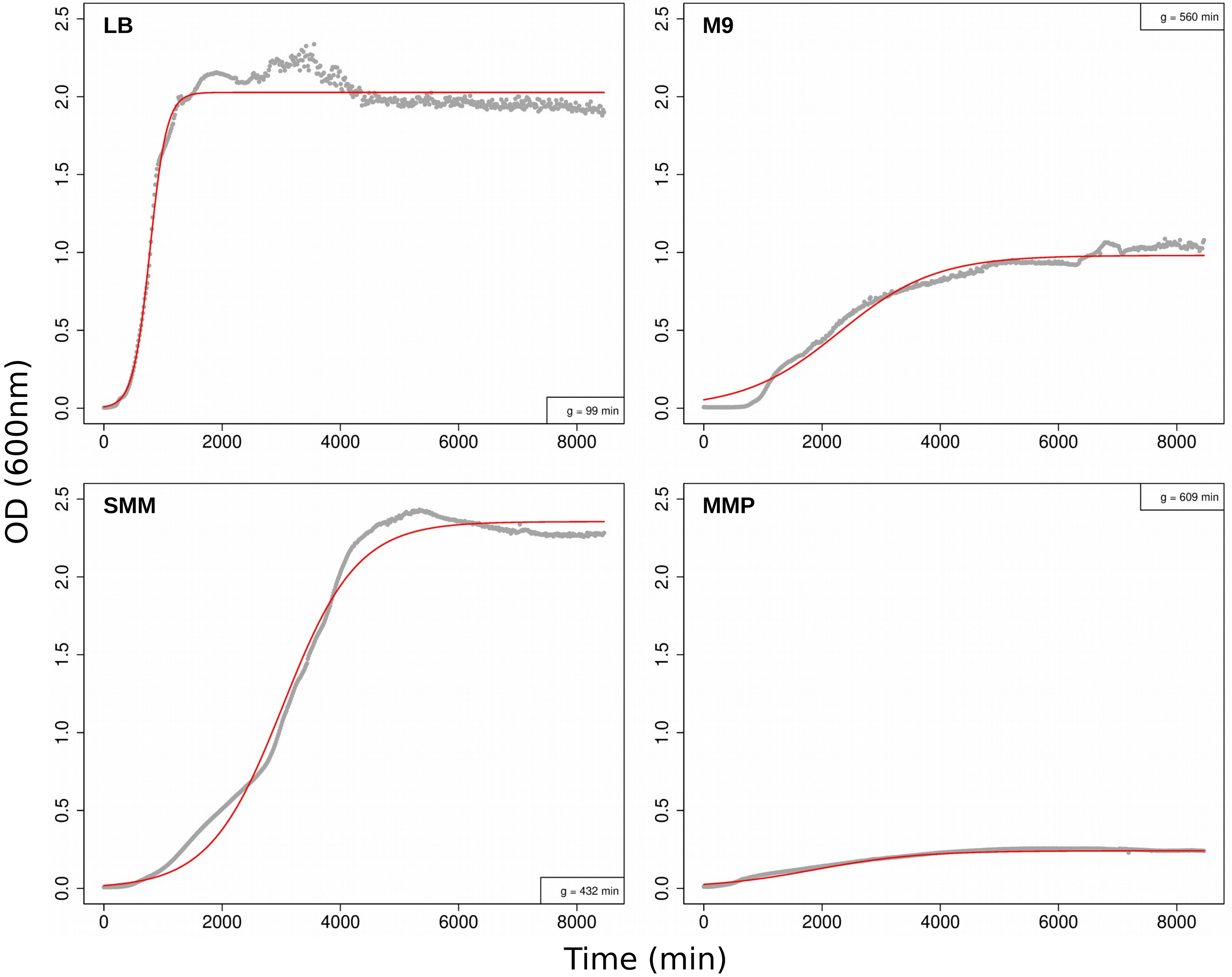

### Supplementary figure S6

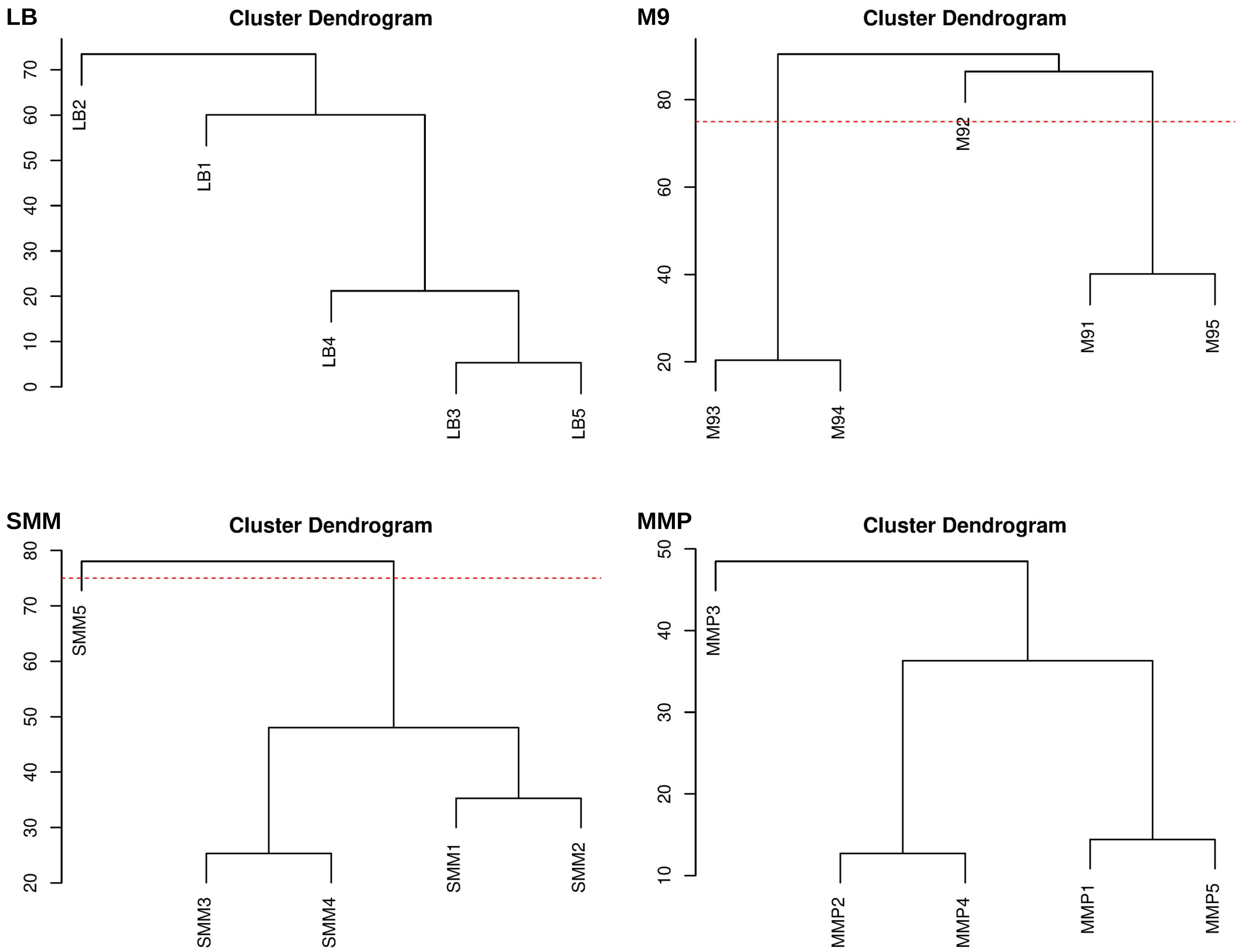

### Supplementary figure S7

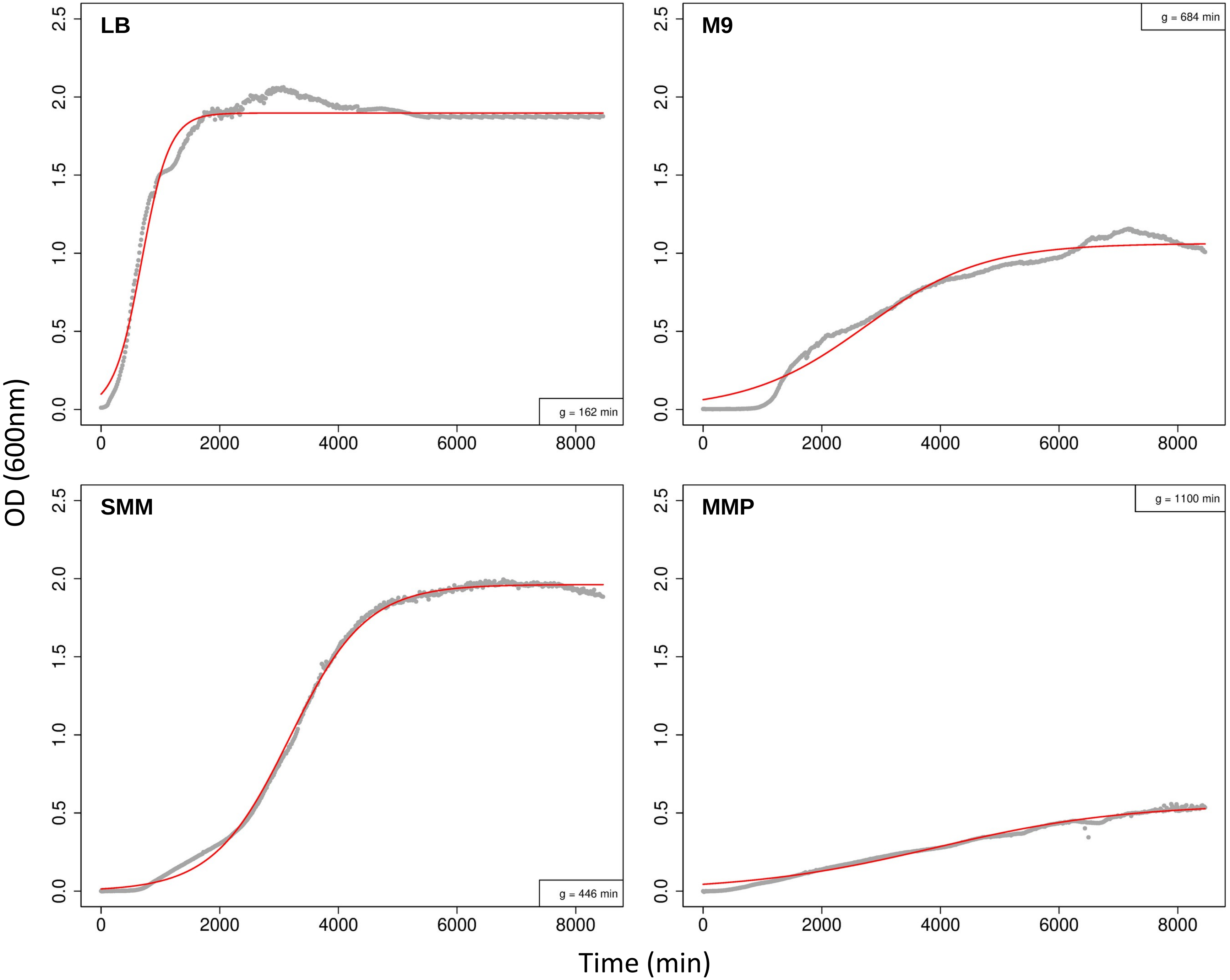
